# Mitochondrial priming in human germ cell tumors is dependent on MCL1 and BCL2L1

**DOI:** 10.64898/2026.09.22.752028

**Authors:** Sirli Anniko, Sonia Khan, Jamie R. J. Morton, Wajiha Iftikhar, Ana A. M. Teixeira, Craig R.G. Willis, Kirsten Riches-Suman, Krzysztof Poterlowicz, Steven D. Shnyder, Rod T. Mitchell, Sherif F. El-Khamisy, Simon J. Allison, Peter K. Nicholls

## Abstract

Germ cell tumors (GCTs) are highly sensitized to cell death in response to DNA damaging agents, a property that underlies the success of current chemotherapeutic regimens. To address the molecular basis for this, known as apoptotic priming, we evaluated how different BCL2 family members modulate the heightened sensitivity of GCTs to therapy. Our analysis of human GCTs finds consistently high expression of the pro-survival factors *MCL1* and *BCL2L1* (*BCLX*) in a cohort of primary tumors and in their embryonic precursor cells, frequently accompanied by copy number gains of these loci and reciprocal losses of their pro-apoptotic interaction partners and inhibitors, *PMAIP1* (*NOXA*) and *BAD*. We find that co-inhibition of MCL1 and BCLX using selective BH3 mimetics results in a potent synthetic lethality in multiple GCT embryonal carcinoma cell lines. When these cell lines were cultured with the DNA damaging agents cisplatin or etoposide, inhibition of MCL1 or BCLX potentiated their apoptotic effect in undifferentiated embryonal carcinoma cell lines, but not in retinoic acid-differentiated cells. The inhibition of MCL1 also heightened cisplatin sensitivity in p53- deficient or -mutant cell lines, which is associated with resistance to therapy. Employing an *in ovo* human xenograft model, we validate that the combination of cisplatin and MCL1 inhibition enhanced the therapeutic response by eliminating tumor cells. Our findings identify MCL1 and BCLX as critical factors to maintain GCT viability and as putative therapeutic targets to further augment GCT responsiveness to DNA damaging agents.

## INTRODUCTION

The remarkable clinical response of germ cell tumors (GCTs) to DNA damaging agents has long been associated with the stem cell-identity and hypomethylated status of tumor cells^1–5^. Pluripotent embryonic stem (ES) cells are similarly highly sensitized to genotoxic stress and are considered poised to undergo mitochondrial-dependent apoptosis, requiring a low level of additional stress to exceed the apoptotic threshold^6^. This sensitivity, known as mitochondrial priming, is dependent on the balance between pro- and anti-apoptotic BCL2 proteins. Functional profiling of human GCTs using BH3 peptides that mimic BH3-only proteins, demonstrates an elevated apoptotic sensitivity relative to adjacent normal tissue^7^. Modelling of these tumors using embryonal carcinoma (EC) cell lines has previously implicated *TP53* (*p53*) and its downstream pro-apoptotic target genes *BBC3* (*PUMA*) and *PMAIP1* (*NOXA*) in the propensity for apoptosis in response to cisplatin^8–10^. This sensitivity is dependent on the stem cell status, as siRNA-mediated knockdown of *POU5F1* and the consequent disruption to the pluripotency network desensitizes GCT cell lines to cisplatin^9^. While the stem cell status and the balance of BCL2 factors direct the propensity for cell death, the pro-survival factors that modulate the apoptotic threshold in GCTs have not been systematically evaluated. Clinical approaches that promote apoptosis by inhibiting pro-survival BCL2 factors could further sensitize tumors to genotoxic agents thereby avoiding toxicity in other tissues, with the potential to enhance the efficiency of existing therapy and overcome resistance in refractory disease^11^.

In this study, we assessed the role of pro-survival members of the BCL2 family as a potential therapeutic target. Using GCT cell lines, we demonstrate that selective inhibition of MCL1, and to a lesser extent BCLX, markedly enhances cisplatin-induced apoptosis in undifferentiated EC cell lines.

## RESULTS

### Testicular germ cell tumors demonstrate genomic variation at key BCL2 family genes

To identify pathways contributing to the sensitivity of germ cell tumors (GCTs) to DNA damage, we examined the expression of the BCL2 family in The Cancer Genome Atlas-Testicular Germ Cell Tumor (TCGA-TGCT) dataset^1^. We identified broad expression of BCL2-related genes, with high expression of *MCL1* (Fig. 1A). By contrast, the BH3-only sensitizers to apoptosis including *BMF*, *BCL2L11* (*BIM*), *PMAIP1* (*NOXA*) and *BAD* showed modest expression in tumors, with universal expression of the apoptotic effector genes *BAX* and *BAK1* (Supplementary Fig. 1A for full dataset).

**Figure 1:**
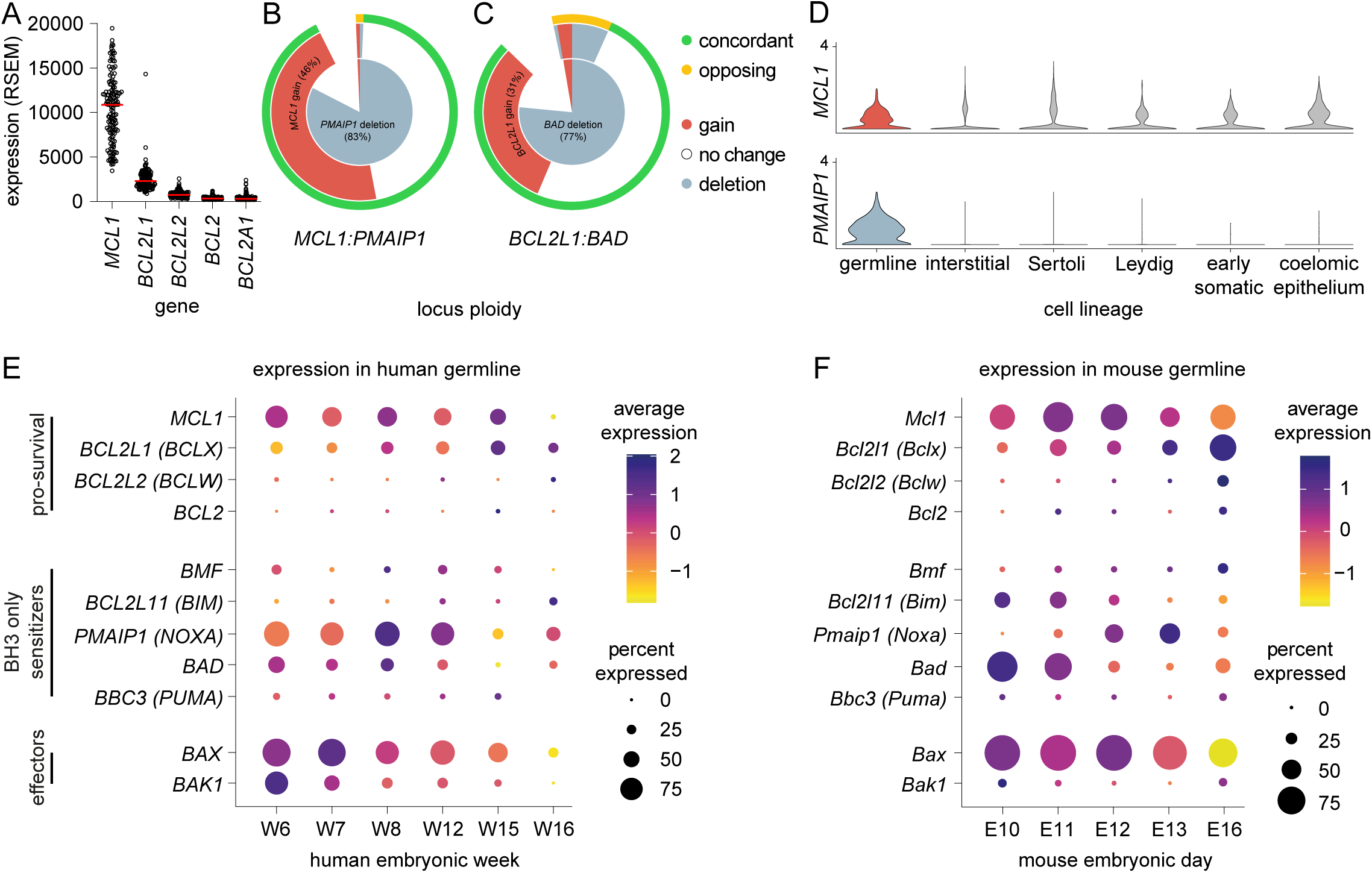
Expression and genetic variation of BCL2-related factors in human germ cell tumors. A) Dot plot of pro-survival BCL2-related factor mRNA expression in human germ cell tumors from the TCGA dataset, ordered by expression level (n=149). Each datapoint represents a single tumor, median expression indicated by red line. B-C) Concentric pie chart of genomic copy number gains and losses of functionally antagonistic BCL2-family pairs. Innermost circle shows deletions (blue) and gains (red) to pro-death genes B) *PMAIP1* (*NOXA*) and C) *BAD*. Second layer shows gains (red) and deletions (blue) to pro-survival factors B) *MCL1* and C) *BCL2L1* (*BCLX*). Outermost circle shows the frequency of functionally concordant gains in the pro-survival and/or gene deletion in the pro-death gene (green) or opposing changes (yellow). Functionally opposing changes (a gain or deletion in both genes of a pair are indicated in yellow). D) Violin plot of single-cell RNAseq analyses of selected genes and lineages in human embryonic gonads. Y-axis presents log-transformed expression of *MCL1* (upper) across multiple lineages and of *PMAIP1* (lower) in the germline. E) Bubble plots showing selected gene expression of BCL2-related factors in embryonic human germline by gestational week (W), and F) mouse germline by embryonic day (E). Color indicates average expression level, circle size reflects percentage of cells expressing gene. See Supplementary Fig. 1-2 for full datasets.

Analyzing genomic copy number variation in the same cohort of tumors demonstrated that 123 (82.6%) contained shallow or deep copy number loss – with one gain – at the locus encoding *PMAIP1*, whose key function is as an endogenous inhibitor of MCL1 (Fig. 1B, inner ring; Supplementary Fig. 1B,C). In parallel with prior reports in GCTs and other cancer contexts^12,13^, the locus encoding the pro-survival factor *MCL1* displayed copy number gains in 69 (46.4%) GCTs, with a single tumor showing copy number loss, indicative of an intolerance to copy number loss (Fig. 1B, middle ring). Collectively, 92.6% of GCTs show functionally concordant gains of *MCL1* and/or losses of *PMAIP1* (Fig. 1B, outer ring). This cohort of GCTs also demonstrated gains of the pro-survival *BCL2L1* locus (*BCLX,* 33.6%) or losses of its inhibitor *BAD* (76.5%), with 80.5% carrying functionally concordant gains and/or losses of this antagonistic pair (Fig. 1C). By contrast, other BCL2 pro-survival factors showed low expression in GCTs, and BH3-only apoptotic sensitizers showed a mix of genomic gains and losses (Supplementary Fig. 1C).

Reciprocal gains of *MCL1* and *BCL2L1*, accompanied by losses of their inhibitors *PMAIP1* and *BAD*, respectively, implies a role for the BCL2-family in the pathogenesis of disease. To assess if this pattern of variation was reflected by expression patterns in the embryonic germline – from which GCTs arise – we assessed the BCL2-family in human and mouse gonadal lineages. In humans, the pro-survival BCL2 factors *MCL1* and to a lesser extent *BCL2L1* (*BCLX*) were broadly expressed among lineages of the embryonic human gonad, whereas *PMAIP1* was predominantly expressed by the germline (Fig. 1D, Supplementary Fig. 1D-F). When analyzing expression patterns within the germline, *MCL1* was the predominant pro-survival factor expressed between week 6 and 16, with lesser expression of *BCL2L1* (Fig. 1E). Among the BH3-only sensitizers, *PMAIP1* was the predominant factor expressed in the newly gonadal germline with expression declining after week 12, a pattern mirrored by the apoptotic effectors *BAX* and *BAK1* (Fig. 1E, Supplementary Fig. 1G).

We next examined if this pattern of embryonic expression was shared in other mammals.

Similarly in mice, *Mcl1* and to a lesser extent *Bcl2l1* were each expressed in the embryonic germline, with *Bax* being the predominant effector (Fig. 1F; Supplementary Fig. 2,3). Like humans, *Pmaip1* was only present in the germline, with transient expression occurring shortly after gonadal colonization when p53 function for programmed cell death is critical to germline fate^14^, while the broad-acting BH3-only sensitizers *Bcl2l11* and *Bad* were present at earlier stages (Fig. 1F, Supplementary Fig. 2).

Given PMAIP1 is a selective inhibitor of MCL1’s pro-survival function, this expression data implies that MCL1 is under additional regulatory control in the embryonic germline during the late migratory and early gonadal stage of development in humans and mice, and additionally, that modulation of this balance – evidenced by concordant genomic alterations in GCTs – may contribute towards cell survival and disease pathogenesis.

### BH3 mimetics sensitize EC cell lines to DNA damaging agents

Cell lines derived from human GCTs are widely used as an *in vitro* surrogate of these tumors. Reflecting their origin, embryonal carcinoma (EC; NT2-D1, 2102EP and NCCIT), yolk sac tumor (GCT72), choriocarcinoma (JAR) and seminoma (TCam-2) cell lines express pro-survival BCL2 genes in a similar pattern to the embryonic germline at gonadal colonization, and of GCTs, with the predominant expression of *MCL1* and expression of BH3-only sensitizers, including *PMAIP1* (Supplementary Fig. 4A). To assess the contribution of the BCL2 family as modulators of apoptosis, we cultured NT2-D1, 2102EP and NCCIT cell lines as models of undifferentiated EC and tested the effect of pharmacologically inhibiting pro-survival factors using selective BH3 mimetics. Once a sublethal concentration for each BH3 mimetic in each cell line was identified (Supplementary Fig. 4), we performed dose response analyses to cisplatin in the presence or absence of a BH3 mimetic (Fig. 2A-C). Correlating with the expression of BCL2 factors by RNAseq, NT2-D1 cells show a modest potentiation of cisplatin by Navitoclax (pan-iBCL2; ABT263, a pan-inhibitor of BCL2, BCL2L1 [BCLX] and BCL2L2 [BCLW], Fig. 2A), an effect mirrored by the selective inhibitor for BCLX (iBCLX, A1155463). Given low expression of *BCL2* and *BCL2A1* in NT2-D1 cells, we conclude that Navitoclax’s potentiation of cisplatin’s effect is predominantly mediated though inhibition of BCLX.

**Figure 2:**
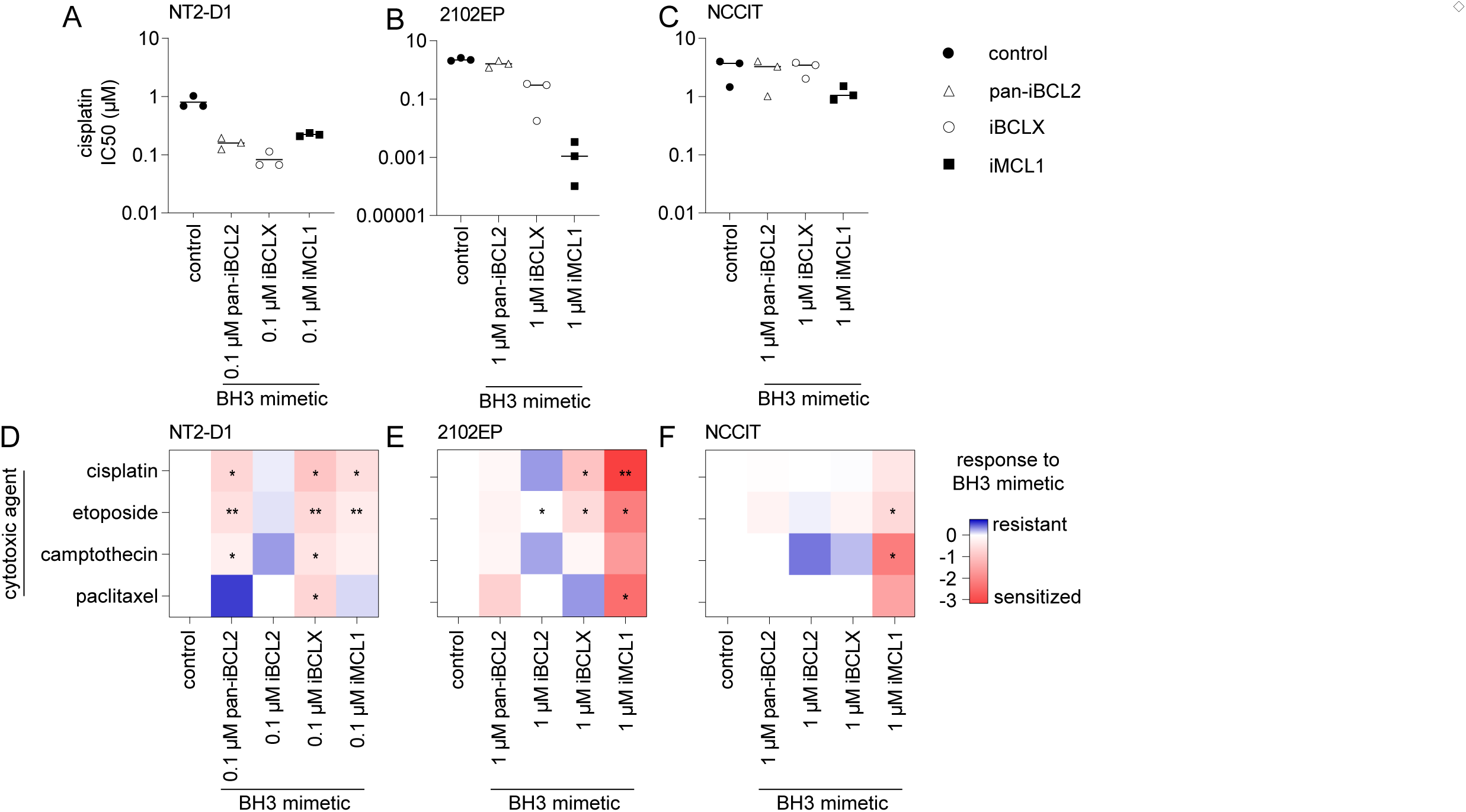
BH3 mimetics sensitize embryonal carcinoma cell lines to cytotoxins. A-C) MTT of EC cell viability in response to fixed sublethal concentration of BH3 mimetic, with the IC_50_ to cisplatin with different treatments. Potentiation by BH3 mimetic is indicated by reduced IC_50_ value. See Supplementary Fig. 4C-H for dose response curves. E-G) Heatmap of potentiation of IC_50_ in EC cell lines with fixed dose of BH3 mimetic (0.1 µM in NT2-D1 cells, and 1 µM in 2102EP and NCCIT cells) with dose-response of cytotoxic agent. Data is log transformed, where value of -1 (red) indicates 10-fold potentiation of cytotoxic effect in the presence of BH3 mimetic. Pan-iBCL2 (ABT263, Navitoclax), iBCL2 (ABT199, Venetoclax), iBCLX (A1155463), iMCL1 (AZD5991). All data normalized to no treatment control, and reflects mean of at least three independent experiments, ± SD. * *p* < 0.05, ** *p* < 0.01

Like BCLX, inhibition of MCL1 (iMCL1, using AZD5991) sensitized NT2-D1 cells to cisplatin (Fig. 2A; p < 0.05). In contrast, 2102EP cells showed only a modest non-significant effect to pan-BCL2 inhibition, while MCL1 inhibition greatly sensitized cells to cisplatin (Fig. 2B; p < 0.01). Unlike either NT2-D1 or 2102EP cells, pan-BCL2 inhibition of cisplatin-treated NCCIT cells showed no potentiation, whereas MCL1 inhibition potentiated cisplatin’s cytotoxic effect (Fig. 2C; *p* = ns). To test if the effect of BH3 mimetics in EC cell lines was unique to the presence of DNA adducts generated by cisplatin, we tested each mimetic in combination with a dose-response to either Etoposide, Camptothecin, or Paclitaxel, which each have distinct mechanisms of action. Each EC cell line demonstrated a broadly similar pattern of response to each of these cytotoxic agents when combined with a BH3 mimetic, indicating that the potentiation observed was not specific to cisplatin (Fig. 2D-F). Notably, potentiation of etoposide by inhibition of MCL1 achieved statistical significance in each cell line, and in two-of-three cell lines for cisplatin. Collectively, these data reveal a critical protective role for MCL1 in each of these GCT cell lines in response to cytotoxic agents. BCLX inhibition potentiated cytotoxic effects in NT2-D1 cells, and to a lesser extent in 2102EP cells, however, its inhibition failed to sensitize NCCIT cells to any of the cytotoxic agents tested (Fig. 2D-F). We additionally considered if BH3 mimetics sensitized other non-EC GCT cell lines to cisplatin. Inhibition of BCLX in both GCT-44 (yolk-sac) and TCam-2 (seminoma) sensitized cells to cisplatin, while inhibition of MCL1 potentiated cisplatin’s effect in both cell lines and additionally in JAR cells (choriocarcinoma, Supplementary Fig. 4I-S).

### Responsiveness to cisplatin and BH3 mimetics in NT2-D1 cells occurs independently of p53

The ability of cells to withstand DNA damage is aided by mutations to *TP53* that preclude a functional response to either initiate apoptosis or to the induction of senescence^15^. Mutation to *TP53* is rare in GCTs compared with other adult cancers^16^, though dominant negative mutations to *TP53* and focal gains of its inhibitor, *MDM2*, have been implicated in resistance to therapy and are associated with poor outcomes^17,18^. We sought to test if depletion of p53 modulates the response to DNA-damage in GCT cell lines, and second, whether selective BH3 mimetics retain their ability to enhance priming for apoptosis. We performed siRNA gene knockdown of *TP53* in p53 wild-type NT2-D1 cells and confirmed efficient knockdown after 48 h by qPCR (Fig. 3A). In NT2-D1 cells, we identified that p53 depletion resulted in a >10-fold reduction in cisplatin’s effectiveness, indicating a functional p53 pathway mediates cisplatin sensitivity in these cells (Fig. 3B), an effect mirrored by the reduced sensitivity of NCCIT cells to cisplatin, which harbor a mutation to *TP53*^19^. The additional pharmacological inhibition of MCL1 using AZD5991 continued to potentiate cisplatin’s effects in p53 knockdown NT2-D1 cells, demonstrating that MCL1 inhibition enhances lethality independent of p53 (Fig. 3B). We additionally explored p53 inhibition and iMCL1 sensitivity in a NT2-D1-resistant subclone^20^. Knockdown of p53 had no effect on cisplatin responsiveness (Fig. 3C), suggesting that an element of this pathway has been altered in this subclone. The pharmacological inhibition of MCL1 in the absence of cisplatin retained some effectiveness but did not impact cisplatin responsiveness in this model with similar IC_50_ values across treatment groups (Fig. 3C). Our data indicates that a functional p53 pathway mediates the hypersensitivity of NT2-D1 cells to cisplatin, and that inhibition of the pro-survival factor MCL1 continues to sensitize EC cells to DNA damage, regardless of p53 status.

**Figure 3:**
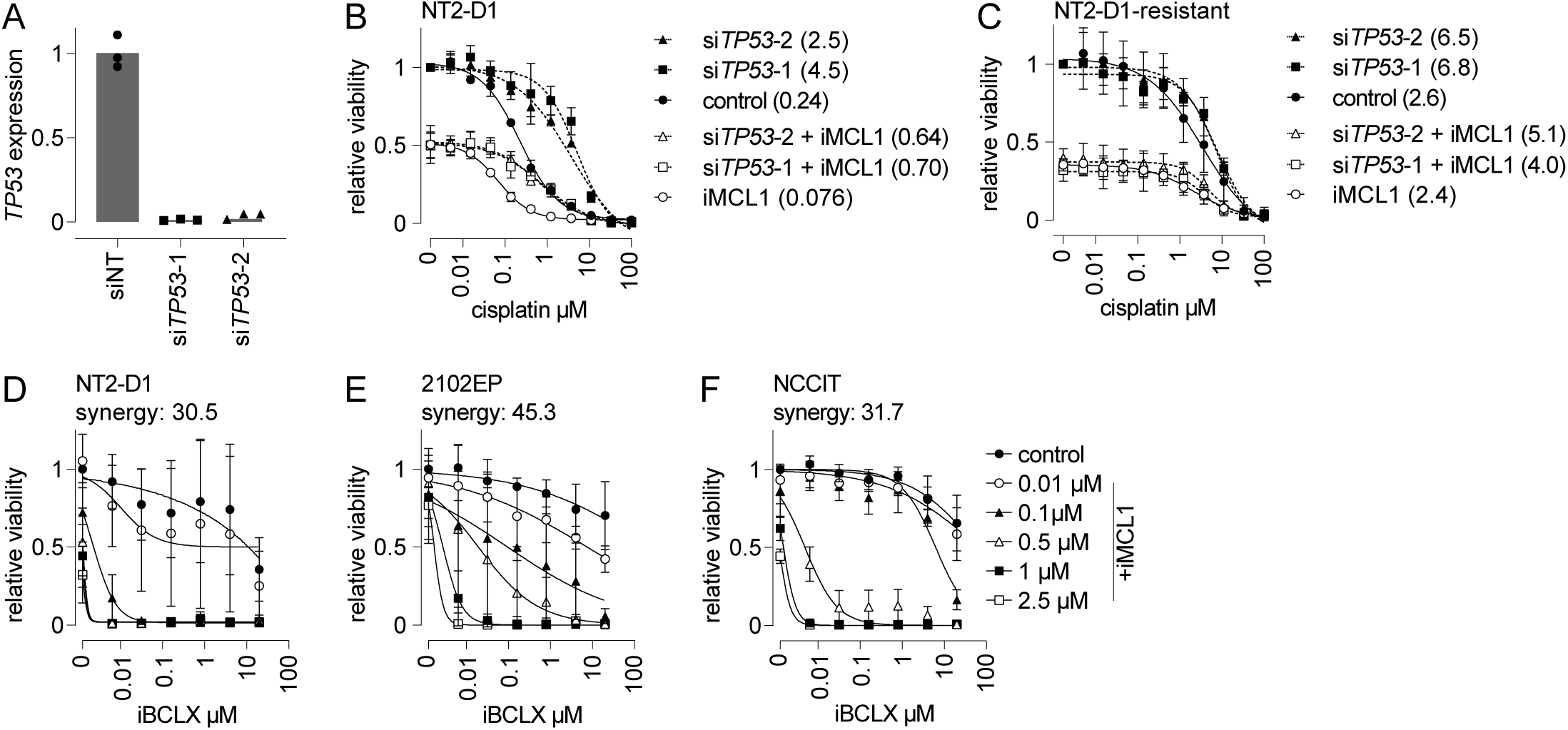
Inhibition of MCL1 potentiates cisplatin independent of p53 status, and is synthetically lethal with BCLX. A) mRNA expression of *TP53* in NT2-D1 cells following normalization to 18S by qPCR after 48 h incubation with a control siRNA (siNT) or with one of two independent p53 siRNAs. Technical replicates from one experiment shown. B-C) MTT assay of cell viability of B) NT2-D1 and C) NT2-D1 cisplatin-resistant subclone. Cells were incubated with siRNA for 48 h, followed by 48 h treatment with control (filled symbols) or with 0.1 µM iMCL1 (open symbols, AZD5991), and cisplatin (x-axis). IC_50_ shown in parentheses beside each condition. Data indicates p53 depletion desensitizes NT2-D1 cells to cisplatin, while iMCL1 continues to potentiate cisplatin. MTT assay of cell viability in D) NT2-D1, E) 2102EP or F) NCCIT cell lines in response to iMCL1 and iBCLX in the absence of cytotoxic agent. Bliss synergy score for each cell line depicted in panel title. Data shows synergism between MCL1 and BCLX in each EC cell lines, with sublethal concentrations of each individual agent yielding lethality when combined with the additional mimetic. All data in panels B-F is mean of at least three independent experiments, ± SD.

### Dual inhibition of MCL1 and BCLX yields synergistic effects in GCT cell lines

Inhibition of either MCL1 or BCLX in NT2-D1 cells demonstrated a modest effect on cell viability in the absence of an additional DNA damage input. To assess if singular-inhibition induces a dependency on an alternative BCL2 factor, we investigated the effect of co-inhibition of both MCL1 and BCLX on GCT cell line viability and analyzed synergism^21^. In each embryonal carcinoma cell line, co-inhibition in the absence of cisplatin yielded complete lethality when cultured with sublethal doses of iMCL1 and iBCLX, indicating that both factors are essential to maintain the viability independent of cisplatin (Fig. 3D-F). The lethal effect in the absence of exogenous DNA damage indicates that MCL1 and BCLX each maintain EC cell viability in conditions where one pro-survival factor is inhibited, confirmed by synergy scores >10 (Fig. 3D-F). The addition of DNA damage overcomes much of this dependency, revealing a greater reliance on MCL1 in each of the EC cell lines assessed.

### Retinoic acid-mediated differentiation modulates BH3 sensitivity

To test if the dependence on specific BCL2 factors for viability in EC cells in response to DNA damaging agents was unique to the undifferentiated status of these cell lines, we differentiated EC cell lines with *all-trans* retinoic acid (RA) and examined their viability in response to BH3 mimetics.

Differentiation was confirmed by the loss of *NANOG* expression indicative of the dismantling of the pluripotency network. After four days in NT2-D1 cells and eight days in NCCIT cells, *NANOG* expression was reduced by >90% (Fig. 4A,F). Within these timeframes, both NT2-D1 and NCCIT cells remained proliferative as indicated by EdU incorporation into DNA being actively synthesized (Fig. 4B,G). RA-differentiated NT2-D1 cells showed a 10-fold reduction in cisplatin sensitivity (p<0.01), but pan-BCL2 or BCLX inhibition continued to potentiate cisplatin’s effect (p<0.01 and p<0.001, respectively), indicating that the effect of BCLX inhibition was not dependent on the stem cell state of the cells (Fig. 4C-D; Supplementary Fig. 5A,B).

**Figure 4:**
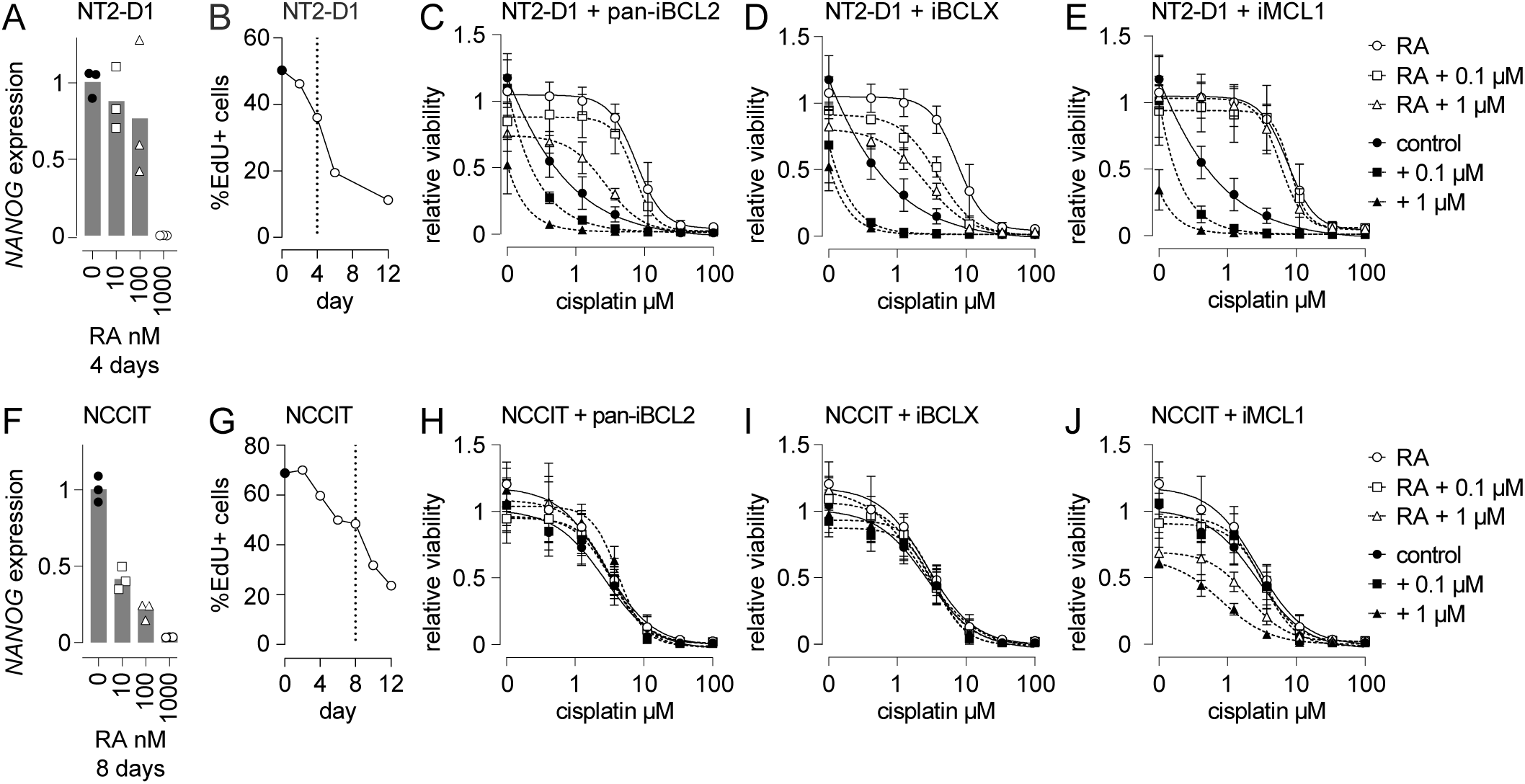
Differentiation of NT2-D1 but not NCCIT cells de-sensitizes to cisplatin and iMCL1. A) Expression of *NANOG* normalized to *18S* by qPCR in NT2-D1 cells after 4 days incubation with *all-trans* retinoic acid (RA, concentration on x-axis). B) Proliferation, initially assessed by DNA synthesis and EdU incorporation across 12 days of 1 µM RA (n=1). C-E) MTT assay of cell viability of NT2-D1 cells treated with 1 µM RA for 4 days (open symbols), followed by 48 h with cisplatin and BH3 mimetic. Data shows RA treatment (open circles) desensitizes cells to cisplatin, which can be partially restored with C) pan-iBCL2 (ABT263, 11.3-fold decrease in viability with 1 µM BH3 vs. RA alone) or D) iBCLX (A1155463, 13.5-fold), whereas RA-treated cells become refractory to E) iMCL1 (AZD5991, 87.5-fold). F) Expression of *NANOG* normalized to *18S* by qPCR in NCCIT cells after 8 days incubation with *all-trans* retinoic acid. G) Proliferation, initially assessed by DNA synthesis and EdU incorporation across 12 days of 1µM RA (n=1). H-J) MTT assay of cell viability of NCCIT cells treated with 1 µM RA for 8 days (open symbols), followed by 48 h with cisplatin and BH3 mimetic. Data shows RA treatment (open circles) has minimal effect on cisplatin sensitivity, and no effect of H) pan-iBCL2 (ABT263) or I) iBCLX (A1155463). RA-treated NCCIT cells show sensitivity to J) iMCL1 at 1 µM, but not at lower concentrations. All data is mean of at least three independent experiments, ± SD. See also Supplementary figure 5 for bar charts depicting relevant statistics.

In contrast, RA-differentiated NT2-D1 cells were completely refractory to MCL1 inhibition, indicating that the dependence on MCL1 was tightly linked to the undifferentiated status of these cells (Fig. 4E, Supplementary Fig. 5C; *p* = ns). Unlike NT2-D1 cells, RA treatment of NCCIT cells had no effect on either cisplatin responsiveness or sensitivity to pan-BCL2 inhibition (*p* = ns), while 1 µM iMCL1 continued to enhance cisplatin’s effect in RA treated cells (Fig. 4H-J; Supplementary Fig. 5D-F, p<0.001).

### In ovo xenograft tumors respond to cisplatin and MCL1 inhibition

As an initial assessment of the anti-tumor efficacy of combining low dose cisplatin with BH3 mimetics and any potential toxicity of this approach, we established *in ovo* GCT tumor xenografts as a preclinical 3Rs ‘proof-of-concept’ model. Human NT2-D1 cells were implanted onto the highly vascularized, extra-embryonic chorioallantoic membrane of fertilized chicken eggs and the effect on tumor growth of iMCL1 in the presence or absence of cisplatin was determined. NT2-D1 cells produced three-dimensional viable tumors with an engraftment rate >90%. These tumors retained an EC histological profile, with dual-labeling of cells with OCT4 and TRA-1-81 (Fig. 5A). To evaluate the efficacy of MCL1 inhibition in this context, tumors were topically treated on E10 with cisplatin, iMCL1, or a combination of both, and effects on tumor weight and embryo viability assessed four days later. Topical administration of cisplatin resulted in a non-statistically significant 25% reduction in tumor mass compared with control (Fig. 5B,C; n=14 each; *p* = 0.155), while administration of iMCL1 alone demonstrated a 16% reduction in tumor mass (Fig. 5C; n=12; *p* = 0.500). By contrast, the addition of both cisplatin and iMCL1 resulted in a 41% reduction in tumor mass (Fig 5C; n=14; *p* = 0.004) although this did not achieve statistical significance from individual treatments. There was no observed adverse effect on embryo viability of the combination treatment compared to control treated tumors (Fig. 5D). Collectively, our data suggests that inhibition of MCL1 potentiates the cytotoxic effects of cisplatin towards embryonal carcinoma cells *in vitro* and *in ovo*, and identifies MCL1 as an important pro-survival factor in the regulation of mitochondrial priming.

**Figure 5:**
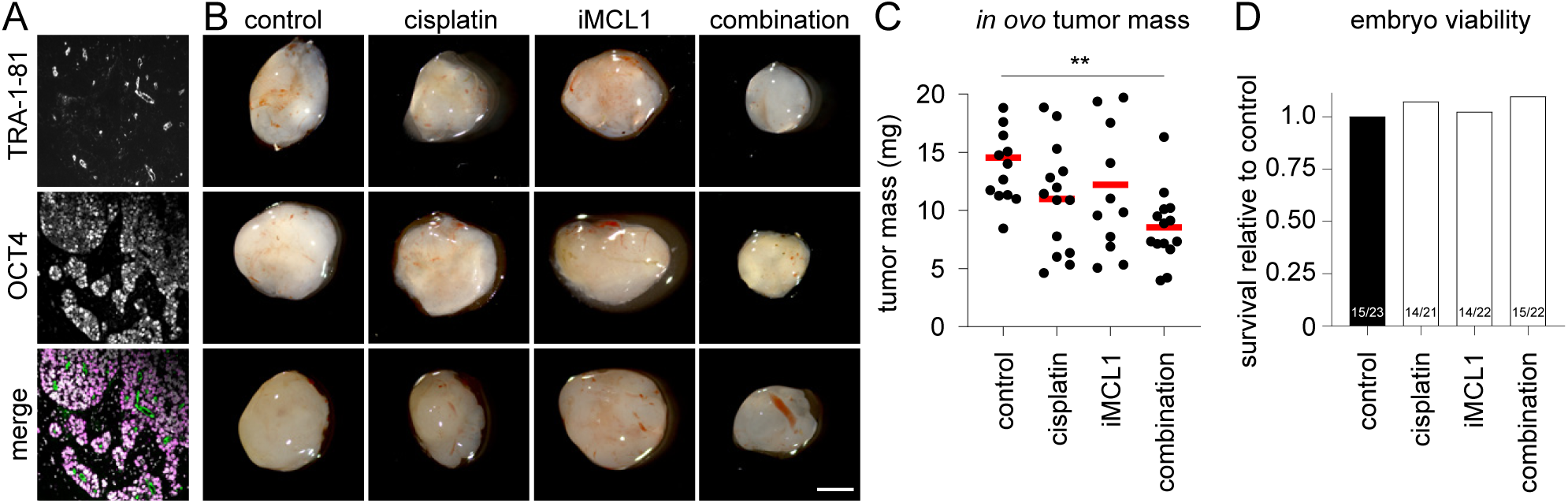
In ovo NT2-D1 tumor xenografts show reduced growth following the combination of cisplatin and iMCL1. A) Immunofluorescence of *in ovo* tumors derived from NT2-D1 cells labelled with TRA-1-81 (upper, green in merge), OCT4 (middle, magenta) and DNA (DAPI, gray in merged lower panel). B) Representative *in ovo* tumors derived from NT2-D1 cells of each treatment group at E14 following topical treatment at E10, scale: 1 mm. C) Mass of tumors following topical administration of cisplatin (10 µM), iMCL1 (AZD5991, 3 µM), or combination (cisplatin + iMCL1) for four days. Data shows individual tumor mass, with red line denoting mean. ** *p* <0.01. D) Chicken embryo viability at E14, normalized to control. Number of surviving embryos indicated at the base of each bar.

## DISCUSSION

### The BCL2 family modulates chemotherapeutic sensitivity in GCT cell lines

Germ cell tumors have long been considered a treatable neoplasm, though the long-term side effects associated with current therapy and the emergence of resistance necessitate novel therapeutic approaches. Using embryonal carcinoma (EC) cell lines as a model of these tumors, we explored the role of the BCL2 family as modulators of the apoptotic threshold in response to chemotherapeutics. Our data reveals a critical role for MCL1, and to a lesser extent BCLX, in maintaining the viability of these cells and as a protective barrier to chemotherapy-induced cell death. When both MCL1 and BCLX were inhibited, EC cells underwent cell death, indicating that both factors sustain cell viability. Consistent with these findings, the *MCL1* locus shows recurrent copy number gains and rare deletions in patient-derived treatment-naïve GCTs^1,12^ suggesting an intolerance to deletion for tumor viability. Mirroring this, its selective inhibitor *PMAIP1* is briefly expressed in the mammalian germline at the time of GCT initiation and shows recurrent copy number losses in >80% of GCTs. In parallel, the *BCL2L1* locus shows frequent copy number gains, while its endogenous inhibitor *BAD* shows copy number losses in most tumors, though their expression is a magnitude lower than *MCL1* and *PMAIP1* in the embryonic germline and tumors. These reciprocal genetic findings in multiple datasets^12^ directly associate the BCL2 family with the survival of GCT cells.

When EC cell lines were treated with a chemotherapeutic agent, such as cisplatin, the simultaneous inhibition of pro-survival BCL2 proteins further sensitized EC cells to DNA damage. In NT2-D1 cells, inhibition of MCL1 or BCLX using a selective BH3 mimetic reduced the concentration of cisplatin required to induce apoptosis. In NCCIT and 2102EP cells, the inhibition of MCL1 alone similarly enhanced the cell death response following DNA damage, an effect similarly observed in TCam-2, GCT-44 and JAR cell lines. Prior analyses have identified GCTs as being primed for apoptosis, and have hypothesized that the BCL2 family mediates the chemosensitivity of tumors to existing therapies^7–9^. Our functional data identify the pro-survival factors MCL1 and BCLX as critical rheostats that modulate apoptotic priming and determine chemotherapeutic sensitivity in human GCT cell lines *in vitro* and *in ovo*.

### Sensitivity to cell death underpins both GCT occurrence and therapeutic outcomes

During mammalian development, primordial germ cells (PGCs) are induced around the time of implantation and later migrate to the genital ridges where they undertake gametogenesis. Supporting

the pivotal role of BCL2 factors to regulate germline viability, deletion of both *Pmaip1* (*Noxa*) and *Bbc3* (*Puma*) in gamma-irradiated mouse embryos enables germline survival beyond lethal checkpoints^22^. By contrast, *Mcl1* deletion in mice disrupts embryo viability, precluding an analysis of its effects in the embryonic germline beyond gastrulation^23^. When cell death is limited via genetic deletion of *p53* or *Bax*, mouse PGCs survive lethal checkpoints and have an increased likelihood of developing GCTs in sensitized mouse backgrounds^24–26^. Similarly in humans, genetic variation at loci associated with the DNA-damage response modulate GCT heritability, including *CHEK2,* and the BCL2-related factors *BAK1*, *BCL2L11 (BAD*) and *BCLAF1*^27–31^. These genetic observations in both mice and humans directly implicate the enhanced survival of germline cells in the initiation and pathogenesis of GCTs^32^.

When GCTs do occur, many remain highly sensitized to DNA damage. The shared pattern of BCL2-family gene expression in the embryonic mammalian germline and in GCTs highlights the mechanisms governing apoptotic sensitivity in these cells. For example, the migratory and early gonadal germline, from which these tumors arise, expresses high levels of the apoptotic cascade including *TP53, PMAIP1, MCL1* and *BAX* in humans and mice. In pluripotent cell lines and in the embryonic germline, this pathway mediates apoptotic priming and cell death^14,33–36^, and *PMAIP1* has additionally been characterized as a factor that modulates long-term fitness of stem cells in culture^37,38^. The temporal regulation of MCL1’s pro-survival function by PMAIP1 in the newly gonadal germline may prime these cells to developmental programmed cell death, such as when the germline fails to undertake commitment^26^ or initiate gametogenesis^14^. This priming of the germline for apoptosis mediated by the BCL2 family^39^ may enable stalled germline cells to be eliminated, thereby limiting their tumorigenic potential.

### Therapeutic potential of BH3 mimetics for germ cell tumors

While GCTs are well treated by the existing therapies of bleomycin-etoposide-cisplatin (BEP), treatment commonly causes long-term side-effects in the comparatively young patient cohort, including an increased risk of secondary cancer as a result of therapy and infertility^40^. A sub-population of individuals remain refractory to existing therapies with loss of p53 tumor-suppressor function one key mechanisms associated with resistance to current therapies^17,18^. The search for new therapeutic approaches is motivated by the need for treatments that are more selective towards tumor cells to decrease side-effects, and that target refractory disease^41^. BH3 mimetics potentially meet both these needs: MCL1 inhibition potentiates DNA damaging agents in undifferentiated EC cell lines with limited effectiveness in differentiated counterparts, which may enable dose de-escalation and the avoidance of BEP-induced systemic side-effects, and second, MCL1 inhibition sensitizes p53-deficient GCT cells that are cisplatin resistant, as seen in other cancer contexts^42–44^. Elevating the feasibility of this approach, *MCL1* is expressed by all GCTs regardless of their histological subtype, with genomic deletions being exceptionally rare. While further validation of this approach is necessary, BH3 mimetics hold potential as a targeted adjuvant therapy to improve the current clinical management of GCTs.

## MATERIALS AND METHODS

### Expression profiling and genomic analysis of TCGA dataset

Gene expression and genomic copy number variation from The Cancer Genome Atlas-Testicular Germ Cell Tumor (TCGA-TGCT, n=149 samples) dataset were obtained via cBioPortal^1,45^. Batch-normalized continuous RSEM values^46^ obtained via cBioPortal were plotted to reflect gene expression levels. Copy number alternations were assessed using the cBioPortal GISTIC algorithm^47^, where a putative gain or shallow deletion were reflected as +1 or -1, respectively.

### Single-cell RNAseq analysis

Human male gonadal samples from post-conception week (PCW) six, seven, eight, 12, 15 and 16 were identified from a published human single-cell RNAseq dataset (GEO accession GSE143356)^48,49^, and mouse gonadal samples from embryonic day (E) 10.5 (described as E10 in text), 11.5, 12.5, 13.5 and 16.5 (GSE184708, Supplementary table 1)^50^, were downloaded from the sequence read archive (SRA). Raw FASTQ files were initially pre-processed using kb-python^51^. Reads were pseudo-aligned to human (GRCh38.p13) or mouse (GRCm39) reference transcriptome, and empty droplets removed using DropletUtils (v1.22.0)^52^ in RStudio. Subsequent analysis was carried out using the standard Seurat (v5.3.0) pipeline in RStudio^53^. Cells expressing >10% mitochondrial genes, <500 features and >150,000 UMI counts in the human and >5% mitochondrial genes, <2000 features and >60,000 UMI counts in the mouse data were excluded from analysis. Doublets were detected using DoubletFinder (v2.0.4)^54^ with the default pN value (25%), resulting in 50,732 human and 139,420 mouse high-quality cells for downstream analysis. Normalization, variable feature identification, scaling and principal component analysis (PCA) were performed using default parameters using Seurat. Fifty principal components were used for community detection by the Louvain method. T-distributed stochastic neighbor embedding (tSNE) was used to visualize cell clusters. Manual cell type annotation using cell-markers identified 15 and 16 cell types in human and mouse, respectively.

### Analysis of BCL2 family expression in human GCT cell lines

To determine expression levels of BCL2-family genes in human GCT cell lines, published RNAseq datasets were reanalyzed (Supplementary Table 3). FASTQ files were downloaded using sratools (v3.1.1). Where present, adapter sequences were removed using Cutadapt (v4.0) and reads of ≥ 35nts retained. Pseudoalignment to the human transcriptome (GRCh38.p13) and transcript quantification were performed with kb-python^51^ (v0.28.2) with the “parity” and “strand” options set according to the library format. Transcript abundances were read into RStudio and converted to gene-level counts using tximport^55^ (v1.38.2).

### Cell culture

Cell lines NCCIT, 2102EP, NT2-D1, NT2-D1-resistant (embryonal carcinoma RRID: CVCL_1451, CVCL_C522, and CVCL_3407, with the resistant subclone previously described^20^), TCam-2 (seminoma, CVCL_T012), GCT-44 (yolk sac, CVCL_A346) and JAR (choriocarcinoma, CVCL_0360) were cultured in RPMI 1640 medium (VWR, 392-0427) supplemented with 10% fetal bovine serum (FBS, Merck, # 7524) and 50 U/mL penicillin-streptomycin (Gibco) at 37 °C and 5% CO_2_. Cells were seeded in a 96-well plate (10,000 cells/well, except JAR: 4,000 cells/well), then treated the next day with combinations of A1155463 (iBCLX, Selleckchem, S7800), ABT-199 (Venetoclax, iBCL2, LKT laboratories, LKT-A0776), ABT-263 (Navitoclax, pan-iBCL2, MedChemExpress, HY-10087), AZD5991 (iMCL1, Cayman Chemical, CAY-28926), cisplatin (MedChemExpress, HY-17394), camptothecin (MedChemExpress, HY-16560), etoposide (MedChemExpress, HY-13629), and/or paclitaxel (MedChemExpress, HY-B0015) for 48 h, as described in the text. Reduction of MTT (Abcam, ab146345) to formazan was taken to reflect cell viability. Cells were incubated with 0.25 mg/mL MTT for 4 h followed by solubilization with DMSO and absorbance measured at 550 nM. Each experiment was performed with four technical replicates, with the number of independent experiments reported in the figure legend.

### Gene expression

Total RNA was extracted using a RNeasy Mini kit (Qiagen, 74104) according to the manufacturer’s protocol, with 1 µg of RNA used for cDNA synthesis using RevertAid First Strand (ThermoFisher Scientific, K1622). Quantitative PCR was performed using Quantstudio 3 (ThermoFisher Scientific) and PowerUp SYBR Green Master Mix (ThermoFisher Scientific, A25776) with 40 cycles of 95 °C for 1 second, 61°C for 20 seconds, 72°C for 15 seconds, followed by a melt curve. cDNA was diluted 20-fold and Ct values were normalized to *18S rRNA* (*RPS18*). Primers are listed in Supplementary Table 2.

### Retinoic acid differentiation of EC cells

*All-trans* retinoic acid (RA) dissolved in water (Thermo-Fisher Scientific, 10552611) was applied to EC cell lines with media replaced every 48 h, and differentiation confirmed measuring *NANOG* relative to *RPS18* by qPCR. To assess cell proliferation, cells were cultured with 10 µM EdU (C10632, Invitrogen) for 2 h. They were then trypsinized, fixed in 4% formaldehyde, washed using Click-It permeabilization reagent, then incubated in the Click-It reaction mix for 30 min, then resuspended in 1% BSA in PBS and analyzed by flow cytometry (Accuri C6 Plus analyzer, BD Biosciences). For MTT assays after differentiation protocols were applied, cells were re-seeded and cultured for a further 48 h with RA and additional pharmacological agents.

### RNA interference

EC cells were transfected by XtremeGene 360 (Roche) using sequences in Supplementary Table 3. siRNA at 25 nM was mixed with XtremeGene 360 in a 3:2 ratio (3 µL siRNA:2 µL transfection reagent) in serum free RPMI-1640 media then added dropwise to cells and incubated for up to 72 h. Knockdowns were confirmed using qPCR.

### In ovo efficacy studies

Fertilized Shaver Brown hen eggs (Medeggs Ltd UK) were used in accordance with the UK Animals Scientific Procedures Act, with embryos humanely terminated on E14. Eggs were incubated at 37.8 °C and 45 % humidity and windowed on E3 as previously described^56^. At E7, eggs containing viable embryos were implanted with 4 x 10^6^ NT2-D1 cells per egg in a 1:1 ratio with Matrigel (4 mg/mL final concentration, Sigma, CLS356237) within a 6 mm silicone ring placed on the chorioallantoic membrane (CAM). On E10, viable eggs were topically treated with 10 µM cisplatin, 3 µM iMCL1, a combination of both, or 0.1 % DMSO solvent control by pipetting 100 µL into the silicone ring containing the implanted tumor cells, and eggs were then incubated a further 4 days. At E14, tumors were excised and imaged using a Zeiss SteREO Discovery V12 stereomicroscope equipped with an Axiocam 305 camera (Zeiss). Attached CAM was removed, tumors weighed and then fixed in 10% neutral buffered formalin for 48 h. Tissues were processed, embedded in paraffin wax, and 5 µm sections prepared. Slides were deparaffinized, rehydrated, then stained with 1:200 dilution of TRA-1-81 (PE conjugated 130-123-334, Miltenyi Biotech) and 1:200 dilution of Rabbit anti-OCT4 (ab19857, Abcam), with OCT4 staining visualized using FITC conjugated Donkey anti-Rabbit IgG (711-095-152, Jackson ImmunoResearch). Slides were mounted with VectorShield containing DAPI, and imaged by fluorescent microscopy. Images were processed in ImageJ (v1.53k).

### Statistics and figure generation

Dose-response curves and absolute IC_50_ values were generated using GraphPad Prism v10.1.1 using non-linear regression (curve fit) with baseline constraint set to 0. A two-way ANOVA test with multiple comparisons (Dunnett) was used to compare IC_50_ values and gene expression by qPCR. Synergism was assessed using the SynergyFinder v3.0 web application using the Bliss model^21^, where scores above 10 denote synergism. A *p* -value < 0.05 was considered statistically significant, unless otherwise indicated.

## ACKNOWLEDGEMENTS

The authors thank Peter Andrews, Daniel Nettersheim and Matthew Murray for cell lines. This work was supported by grant SBF/008/1051 Springboard Award from the Academy of Medical Sciences to PKN, and a Children with Cancer UK grant CwCPG25\100026 to PKN, SDS, RTM and SJA. SA was the recipient of a PhD fellowship from the Faculty of Life Sciences at the University of Bradford. *In ovo* studies were additionally supported by a URF award from the University of Huddersfield to SJA. The authors acknowledge the use of the University of Bradford High Performance Computing Service in the completion of this work.

## AUTHOR CONTRIBUTION

Conceptualization, Writing – Original draft: SA, PKN. Formal analysis and investigation: SA, SK, JRJM, WI, AAMT, SJA, PKN. Funding acquisition and project administration: PKN, SJA, SDS, RTM. Methodology: SA, SK, AAMT. Software: SK, CRGW. Supervision: CRGW, KRS, KP, SEK, SJA, PKN. Visualization: SA, SK, PKN. Writing – review & editing: all authors

## ETHICS

This study did not require ethical approval.

## AVAILABILITY OF DATA AND MATERIALS

All data and materials are available on request.

## SUPPLEMENTARY FIGURE LEGENDS

**Supplementary Figure 1:**
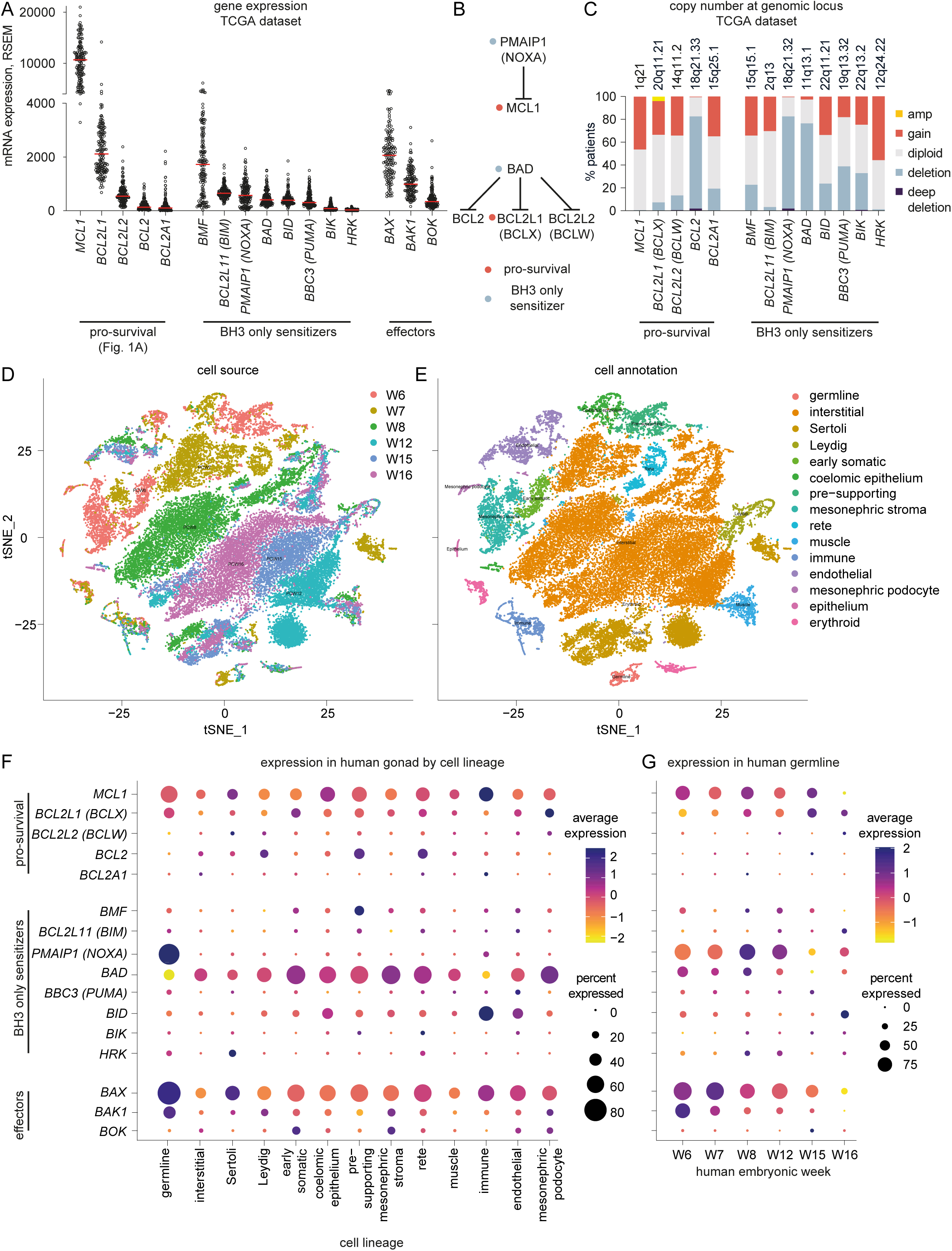
Expression and genetic variation of BCL2-related factors in human germ cell tumors and expression in human embryonic germline. A) Dot plot of mRNA expression of BCL2-related factors expression in human germ cell tumors from the TCGA dataset. Each dot reflects expression in a single tumor, red line shows median expression, n=149. Gene order reflects functional grouping, followed by expression rank. B) Summary diagram of BH3-only pro-death factors PMAIP1 and BAD, and their endogenous pro-survival target. C) Bar chart of copy number variation of pro-survival and BH3-only sensitizers of BCL2 factors in the TCGA germ cell tumor dataset. Deep deletions (purple), deletion (blue), diploid (gray), gain (red) and amplification (amp, yellow), n=149. Cytogenic location shown above each bar; note that BCL2 and PMAIP1 are linked on Chr. 18. D,E) tSNE plots showing human cell clusters colored by D) week of embryonic development, and E) cell annotation based on marker gene expression (see also Supplementary Fig. 3, Supplementary Table 3). All samples are XY (male) as outlined in Supplementary Table 1. F-G) Bubble plots showing gene expression of BCL2-related factors in F) human gonadal cell types identified by tSNE cluster, and in G) embryonic human germline by post-conception week (W). Color indicates average expression level, circle size reflects percentage of cells expressing the indicated gene. Gene order corresponding to Fig. 1A. Data shows high expression of MCL1 (and to a lesser extent BCL2L1) in the human embryonic germline and other gonadal lineages. The apoptotic BH3-only sensitizer PMAIP1 is predominantly expressed in the human germline, whereas BAD is broadly expressed in other gonadal lineages.

**Supplementary Figure 2:**
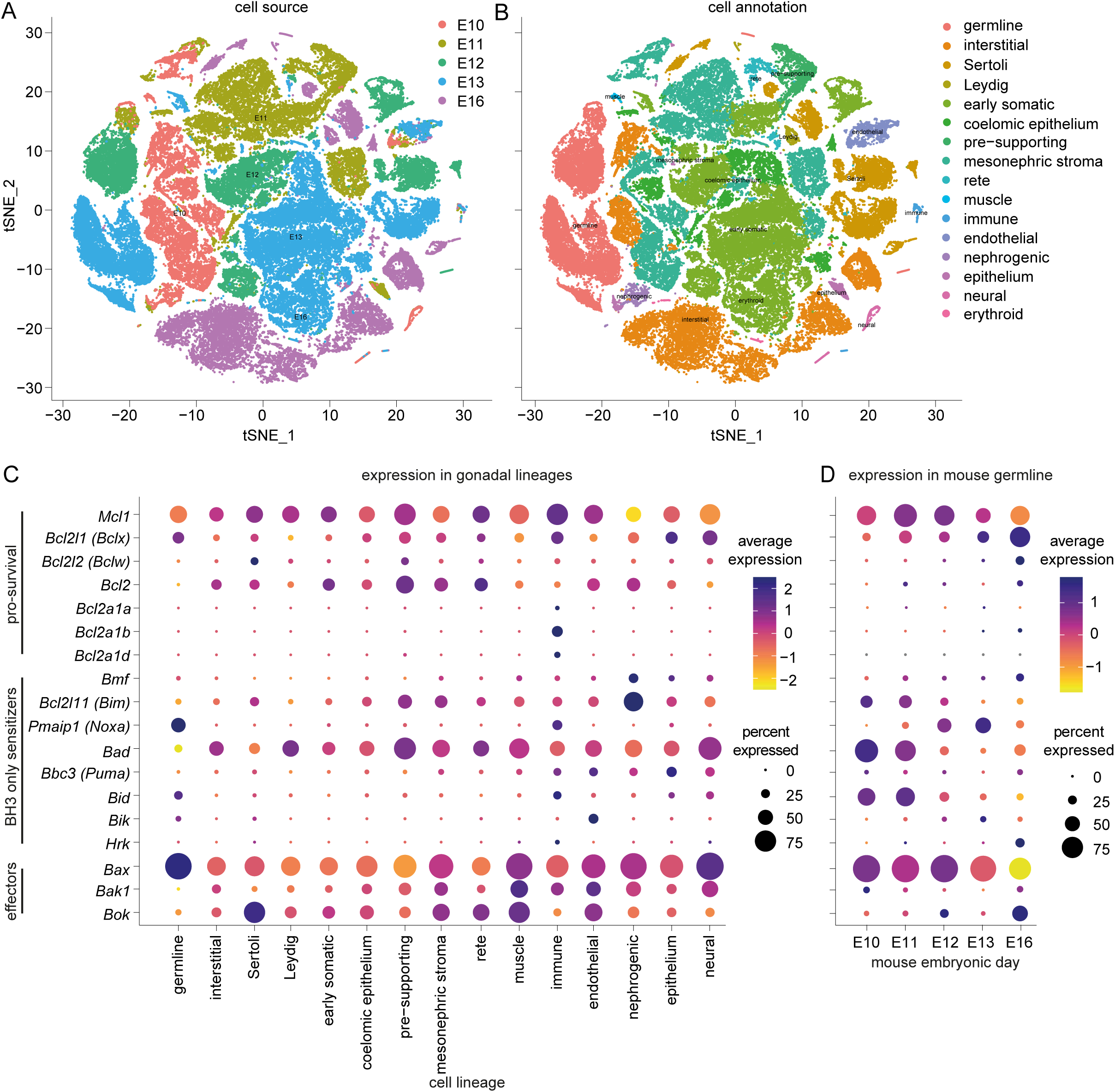
Expression of BCL2-related factors in the mouse embryonic germline. A-B) tSNE plots showing mouse cell clusters colored by A) day of development, and B) cell annotation based on marker gene expression of XY samples (see also Supplementary Fig. 3, Supplementary Table 3). C-D) Bubble plots showing gene expression of BCL2-related factors C) mouse gonadal cell type identified by cluster, and D) in embryonic mouse germline by embryonic day. Color indicates average expression level, circle size reflects percentage of cells expressing gene. Gene order corresponding to Fig. 1A. Data shows high expression of Mcl1 (and to a lesser extent Bcl2l1) in the mouse embryonic germline and other gonadal lineages. The apoptotic BH3-only sensitizer Pmaip1 is predominantly expressed in the germline, whereas Bad is broadly expressed in other gonadal lineages.

**Supplementary Figure 3:**
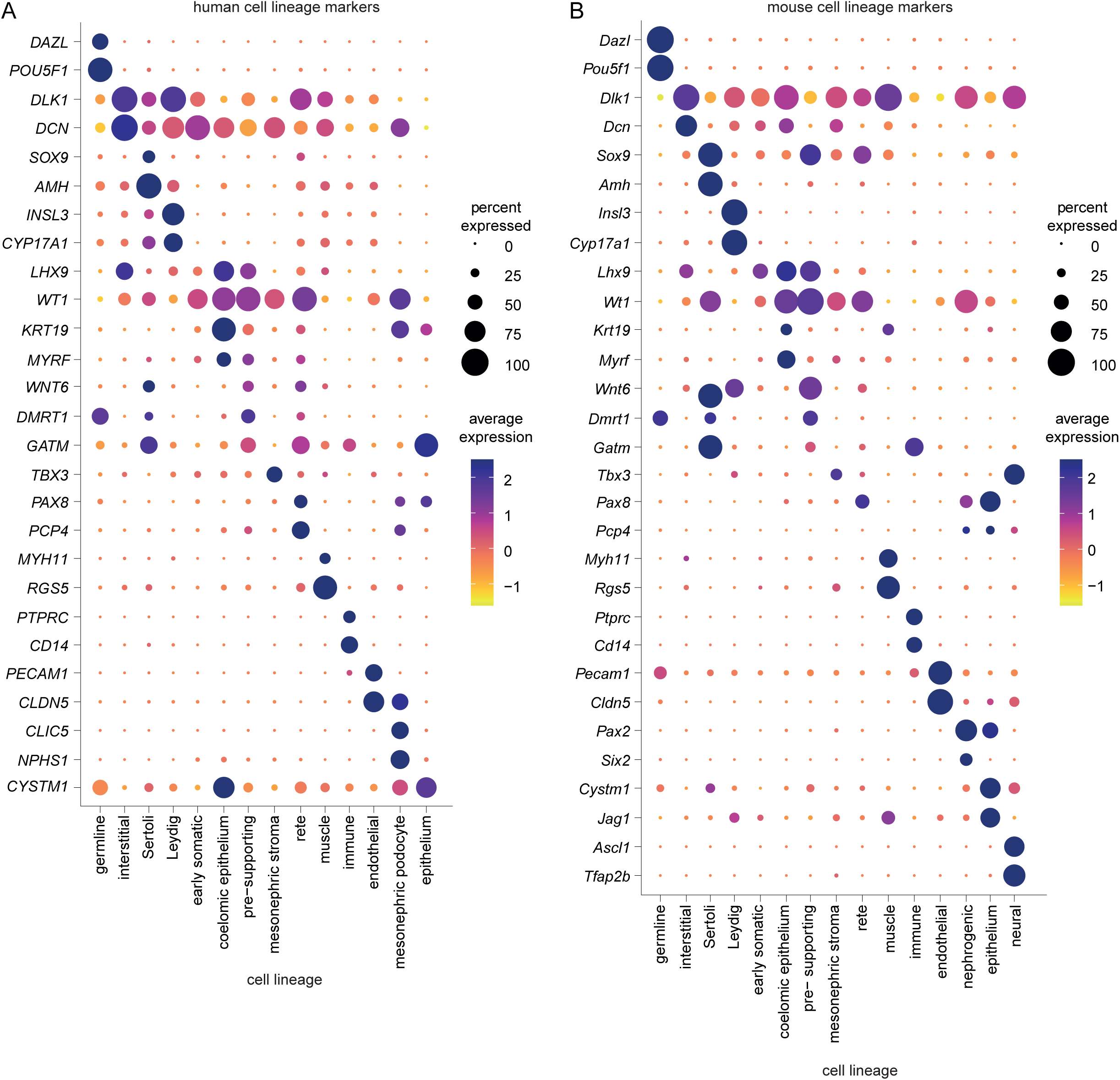
Expression of selected marker genes in identified tSNE plot cell clusters. Bubble plots showing gene expression of selected marker genes for each cell type identified by cluster in A) human and B) mouse. Color indicates average expression level, circle size reflects percentage of cells expressing gene. Gene order corresponding to cluster organization from Supplementary Figures 1 and 2.

**Supplementary Figure 4:**
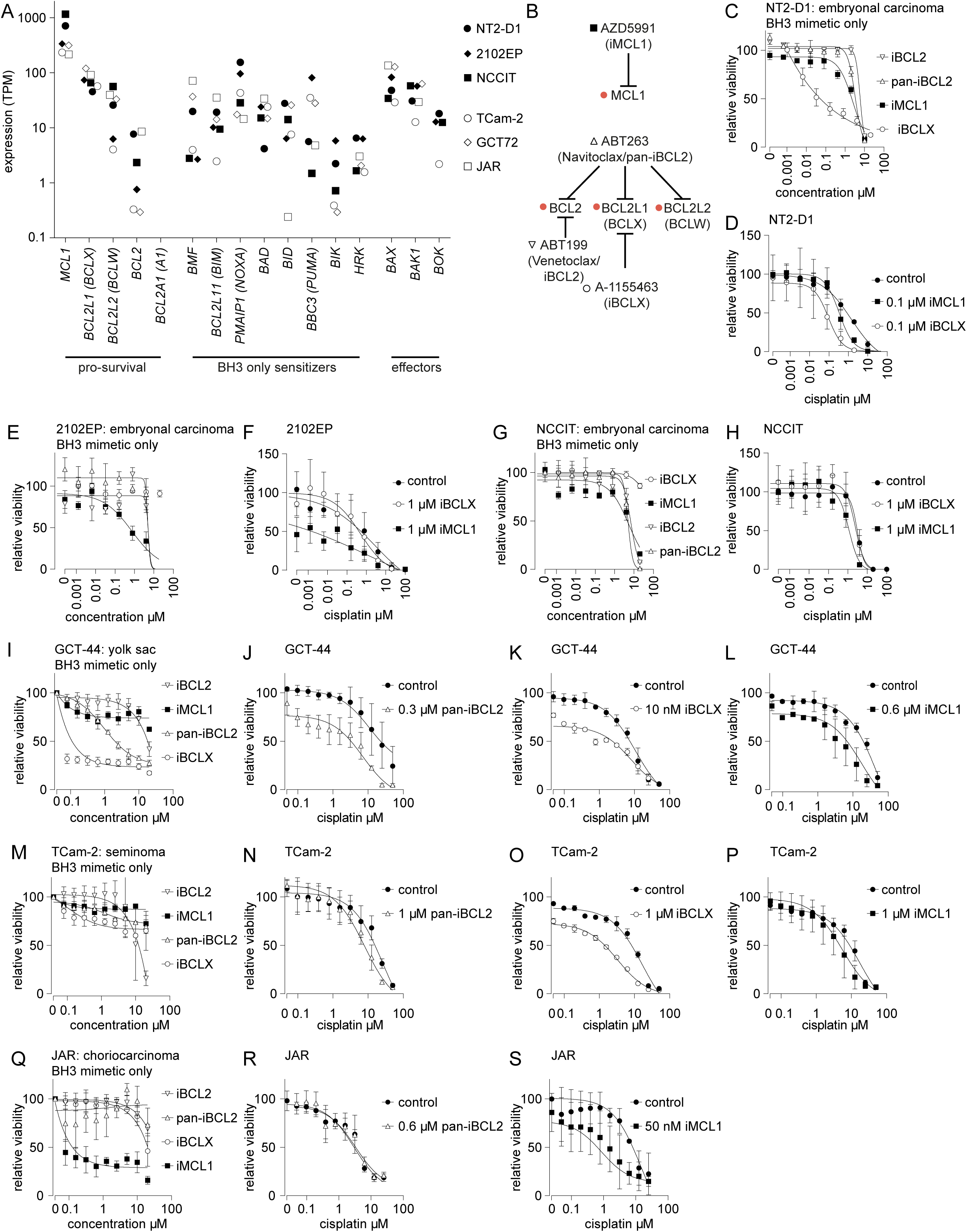
Expression of BCL2 factors in tumors and cell lines and evaluation of BH3 mimetics. A) Dot plot of BCL2-related factor expression in selected human germ cell tumor cell lines by RNAseq, with embryonal carcinoma cell lines shown using filled symbols. Y axis shows mean expression in Transcripts Per Million (TPM). Data points < 0.1 TPM not shown. B) Summary diagram of BH3 mimetic selectivity for specific pro-survival BCL2 factors. C-S) MTT assay of GCT cell line viability in response to pan-inhibition of BCL2 factors (pan-iBCL2, ABT263, Navitoclax), iBCL2 (ABT199, Venetoclax), iBCLX (A1155463) or iMCL1 (AZD5991). C) NT2-D1 cells show sensitivity of BCLX inhibition alone, while D) in combination with cisplatin, inhibition of MCL1 yielded a 3.6-fold potentiation, while inhibition of BCLX produced a 9.7-fold potentiation. E) 2102EP cells showed sensitivity to MCL1 inhibition alone, while F) in combination with cisplatin, inhibition of MCL1 yielded > 1000-fold potentiation, while inhibition of BCLX produced a 10.6-fold potentiation. G) NCCIT cells show minimal sensitivity to BH3 mimetics alone, while H) in combination with cisplatin, inhibition of MCL1 yielded a 2.7-fold potentiation, while inhibition of BCLX produced no effect. I) GCT-44 cells showed sensitivity to BCLX inhibition alone, resulting in 10 nM used in subsequent assays. D) Pan-iBCL2 yielded a 1.5-fold additive effect with cisplatin, which could be replicated with K) iBCLX yielding a 1.3-fold effect. L) iMCL1 yielded a 3.1-fold sensitization to cisplatin. M) TCam-2 cells showed limited sensitivity to BH3 mimetics, with N) 3.4-fold sensitization to cisplatin with pan-iBCL2, while O) iBCLX showed a 5.1-fold potentiation of cisplatin. P) iMCL1 showed a 3.1-fold potentiation to cisplatin. Q) JAR cells showed sensitivity to iMCL1 alone, resulting in 50 nM being used in subsequent assays. R) Pan-iBCL2 yielded no additive effects with cisplatin (meaning that the effect of iBCLX was not investigated), while S) iMCL1 showed a 16.5-fold potentiation of cisplatin. All MTT data normalized to relevant control, and reflects mean of at least three independent experiments, ± SD. * p < 0.05, ** p < 0.01

**Supplementary Figure 5:**
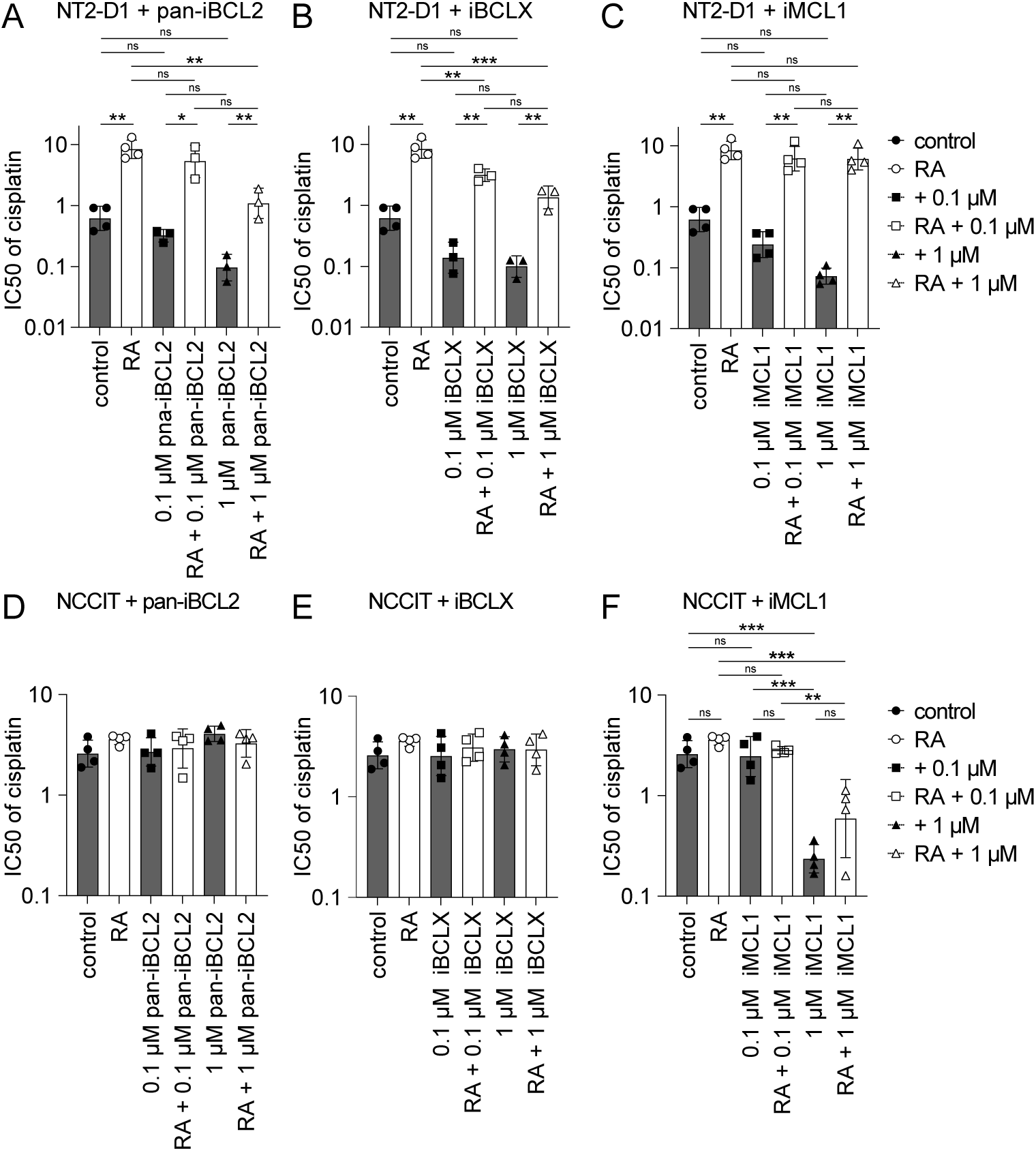
Viability of NT2-D1 and NCCIT cells following differentiation with RA and treatment with cisplatin and BH3 mimetics. IC_50_ values derived from Figure 4 data, with relevant statistical comparisons. RA: 1 µM all-trans retinoic acid, pan-iBCL2: Navitoclax, iBCLX: A1155463, iMCL1: AZD5991. Mean ± SD, ns = not significant, * p < 0.05, ** p < 0.01, *** p < 0.001.

## SUPPLEMENTARY TABLES

**Supplementary table 1:**
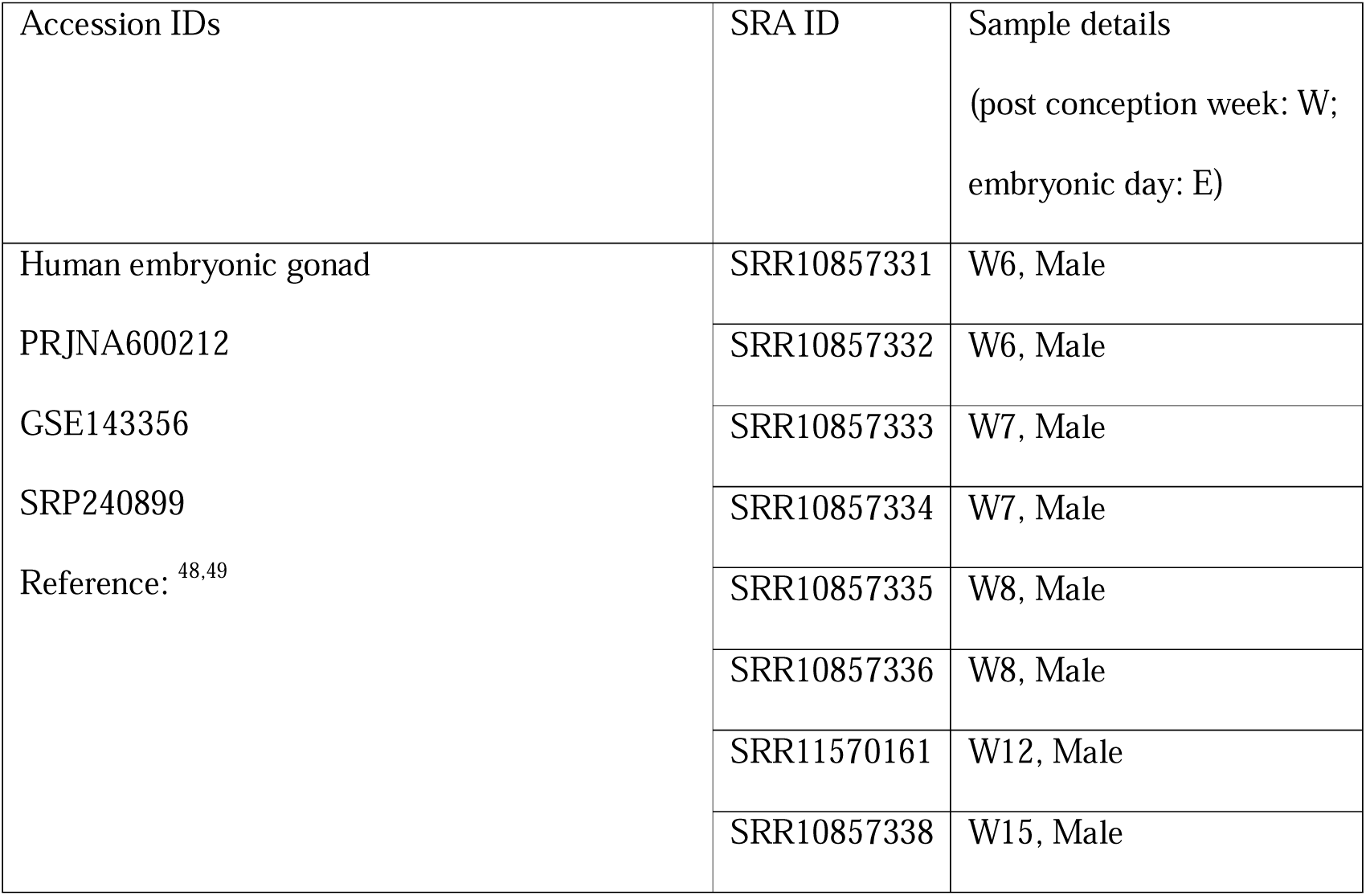
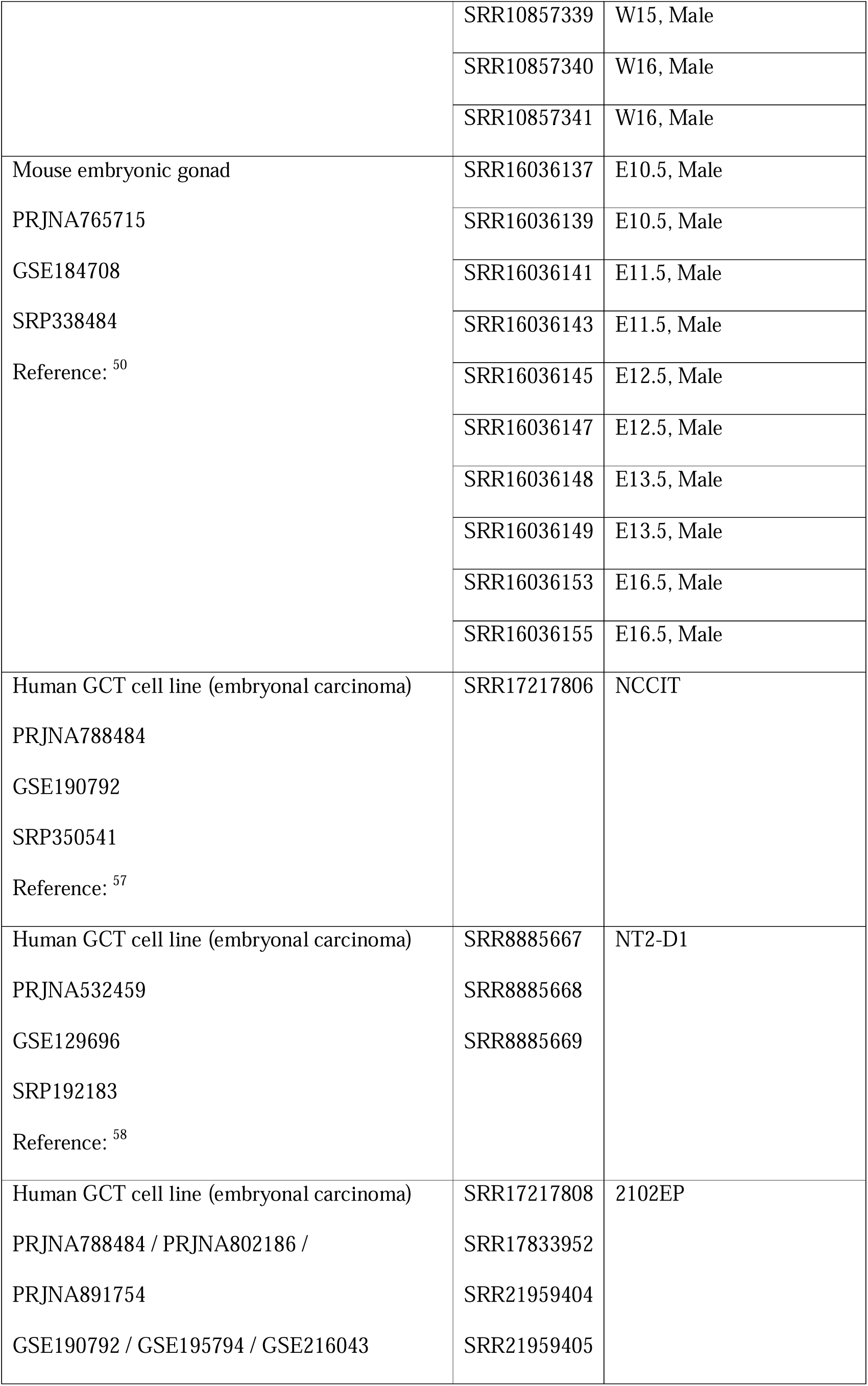
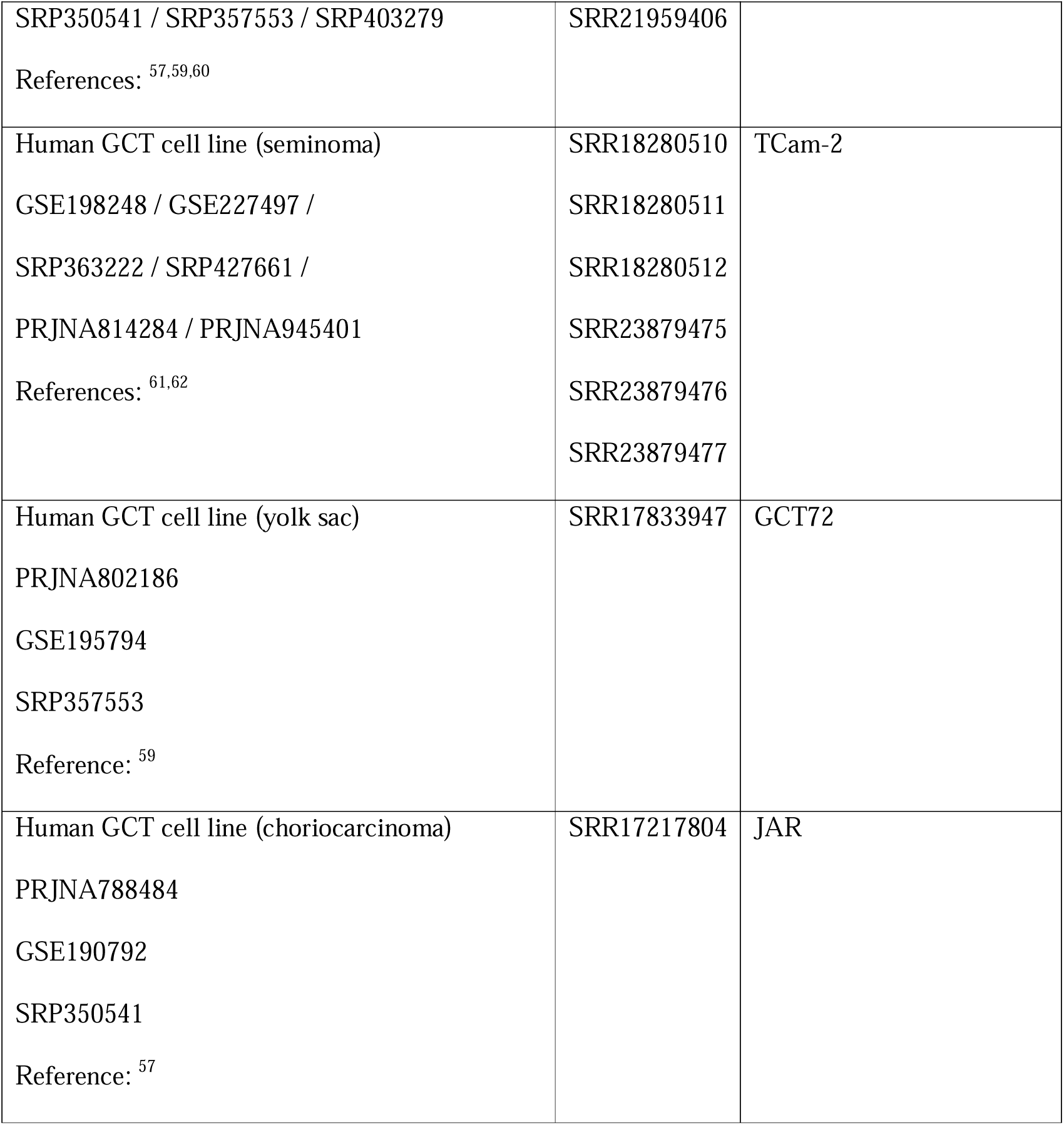
Sample IDs for human and mouse scRNAseq and RNAseq analysis.

**Supplementary table 2:**
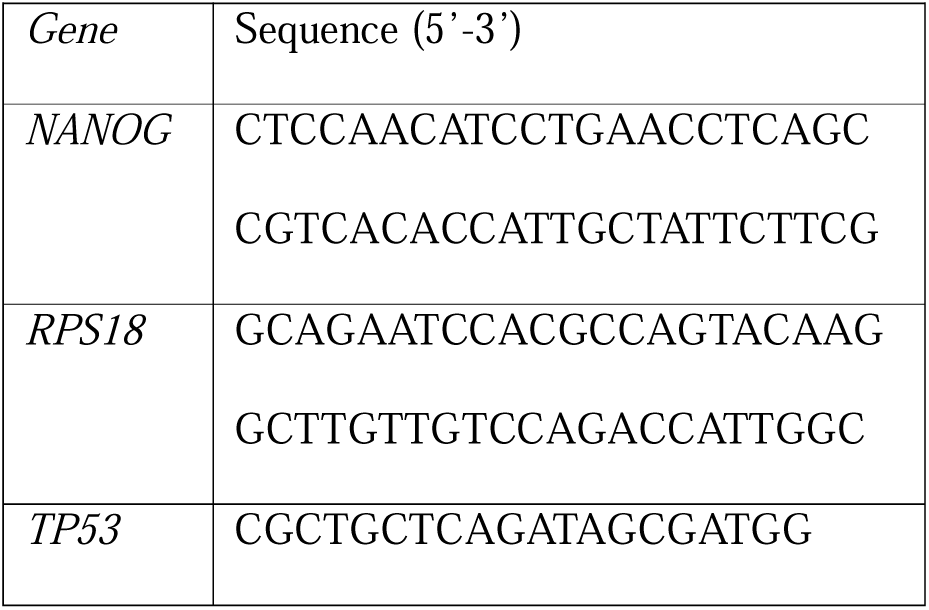
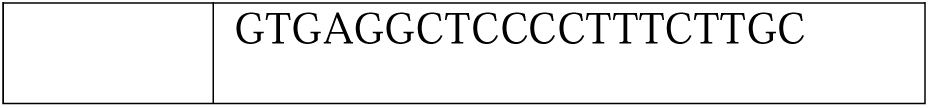
Primers sequences used for qPCR.

**Supplementary table 3:**
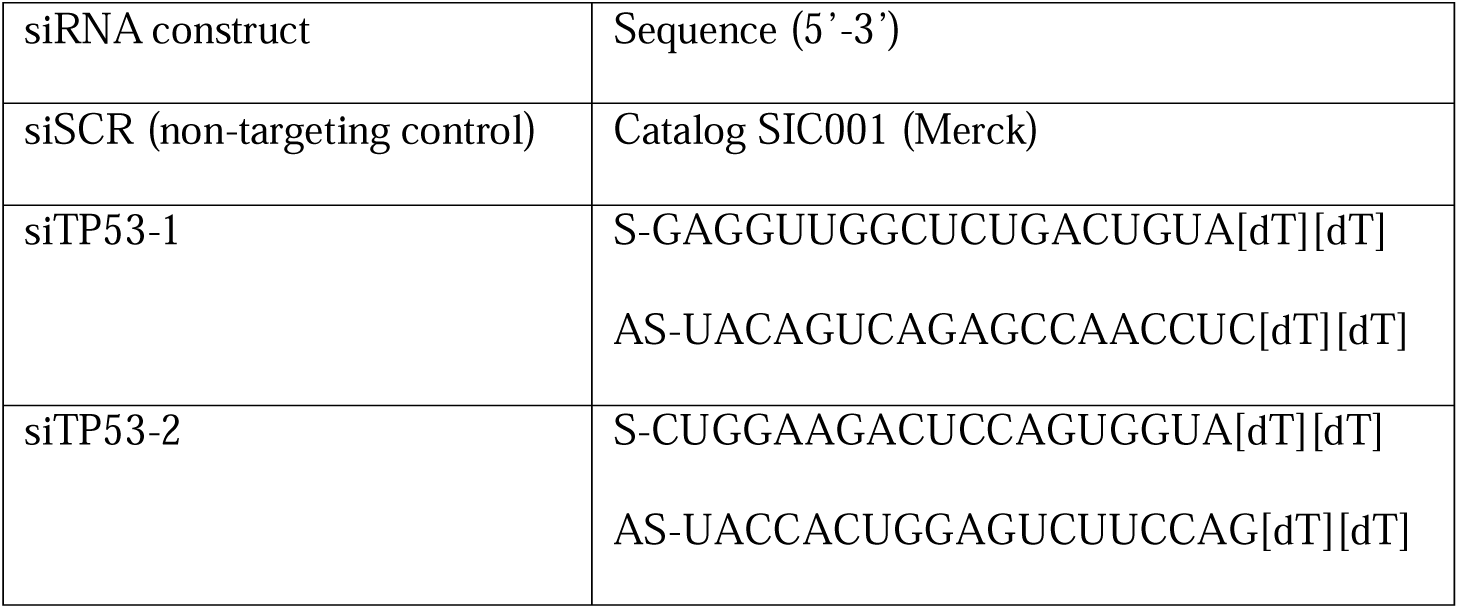
siRNA sequences used for gene knockdown.

